# Screening of FDA-approved small molecules to discover inhibitors of the *Pseudomonas aeruginosa* quorum-sensing enzyme, PqsE

**DOI:** 10.1101/2025.07.25.666791

**Authors:** Hannah A. Jones, Mary J. Baxter, Nicolas Zimmermann, Ada Li, Katelynn A. Perrault Uptmor, Isabelle R. Taylor

## Abstract

*Pseudomonas aeruginosa* is a notorious pathogen that is a leading cause of hospital-acquired infections. Due to its heightened resistance to a broad range of antibiotics, there is great need for new antimicrobial agents that are effective in treating *P. aeruginosa* infections. One attractive option for exploring new antibiotic targets is the quorum sensing (QS) pathway, as it governs many of the pathogenic behaviors that allow *P. aeruginosa* to stage and maintain infections. Within the QS pathway, there is a key protein-protein interaction between an enzyme, PqsE, and one of the master QS regulators, RhlR. Although its catalytic function is dispensable for its interaction with RhlR, previous mutagenic work characterizing the active site of PqsE identified active site mutations that induce a conformational change in PqsE, preventing it from forming a complex with RhlR. These active site mutations, when introduced stably into the genome of *P. aeruginosa*, also lead to a significant decrease in production of a key toxin, pyocyanin, and prevent colonization in the lungs of a murine host. These findings encourage drug discovery efforts to identify small molecules that bind in the active site of PqsE and induce a conformational change similar to the ones induced by particular amino acid substitutions in the active site. Here, we performed a fluorescence polarization screen to identify molecules that bind in the active site of PqsE. We screened a library of FDA-approved drugs, with the intention of potentially repurposing a known molecule for the treatment of *P. aeruginosa* infections. Three molecules were identified, two of which showed inhibitory activity that was consistent with a competitive mode of inhibition. One hit molecule, Apomorphine, had a distinctly different inhibitory profile, and is potentially binding outside of the active site to allosterically inhibit enzyme activity of PqsE. All three hit molecules were tested in an *in vivo* enzyme activity assay, and one of the competitive inhibitors, Vorinostat, was found to inhibit intracellular PqsE. Vorinostat is now being explored as a candidate for synthetic derivatization to inhibit the PqsE-RhlR protein-protein interaction via binding in the PqsE active site.

## Introduction

The human pathogen, *Pseudomonas aeruginosa*, is a leading cause of nosocomial infections and a significant burden to human health ^1,2^. In addition to the many virulence factors produced by *P. aeruginosa*, the species is capable of carrying out coordinated group behaviors such as biofilm formation that contribute to its heightened pathogenicity ^3^. These behaviors are largely under the control of the bacterial cell-to-cell communication system, called quorum sensing (QS). Through the production, release, and detection of small molecule signals, called autoinducers, *P. aeruginosa* uses QS in order to make a coordinated lifestyle switch between carrying out single-cell and group behaviors ^4,5^. This lifestyle switch, which is marked by widespread changes in gene expression, occurs once the concentration of autoinducers in the surrounding environment reaches a critical threshold, signifying the local population has reached high-cell density. Activation of the QS system at high-cell density is critical for the ability of *P. aeruginosa* to stage an infection in a host, and therefore, is of great interest as a target system for the development of new antimicrobial agents ^6^.

There are three main systems that make up the QS network in *P. aeruginosa*, two of which (the Las and Rhl systems) consist of a LuxI/R autoinducer synthase/receptor pair and one (the Pqs system), more specialized to *Pseudomonads*, that consists of a LysR receptor and a biosynthetic operon dedicated to the production of its respective autoinducer ^7^. The Las system consists of a synthase, LasI, responsible for producing the autoinducer N-(3-oxo-dodecanoyl)-L-homoserine lactone (3-oxo-C12-HSL), and the receptor that binds 3-oxo-C12-HSL, LasR ^8^. For the Rhl system, RhlI is the synthase that produces the autoinducer, N-butanoyl-L-homoserine lactone (C4-HSL), which binds the receptor, RhlR ^9^. These receptors, when bound to their respective autoinducers, are activated as transcription factors that control the expression of hundreds of genes involved in group behaviors ^10^. For the Pqs system, the enzymes PqsA-D along with PqsH are collectively responsible for the synthesis of the *Pseudomonas* Quinolone Signal (PQS)^11^ which binds and activates the receptor PqsR. These three systems are highly interconnected, and the regulons of LasR, RhlR, and PqsR consist of genes regulated by all three, two, or only one of the receptors ^12–14^. As an added layer of connectivity, the fifth gene of the *pqsABCDE* operon is dispensable for the production of PQS ^11^, but indispensable for the activation of a large portion of the RhlR regulon ^15–17^. This is due to a non-enzymatic function of the PqsE protein, which is its ability to form a complex with RhlR ^18–20^.

The PqsE-RhlR protein-protein interaction is crucial for the ability of *P. aeruginosa* to carry out a range of behaviors, including the production of a toxin, pyocyanin ^21,22^. Interestingly, mutations introduced to the PqsE active site have been shown to cause a conformational shift that disrupts the interaction with RhlR at a distal site. Specifically, the substitution of an active site glutamate with tryptophan (E182W) causes a loop rearrangement which drastically weakens the affinity of PqsE for RhlR and almost entirely inhibits pyocyanin production ^21,23^. The introduction of the E182W substitution to the PqsE active site was even enough of an alteration to prevent colonization of *P. aeruginosa* in the lungs of a murine host ^23^. For this reason, the PqsE active site is being explored as a target for the development of small molecule modulators, despite the finding that PqsE enzyme activity is dispensable for the PqsE-RhlR interaction ^21^. In this study, we carried out a screen for molecules that compete with an active site-targeted probe, BB562, to bind PqsE. This screen produced molecules that take on different binding modes, and can serve as starting points for the development of PqsE-RhlR allosteric inhibitors, as well as tool compounds for the further characterization of PqsE enzymatic activity. While none of the identified inhibitors were found to affect the PqsE-RhlR interaction, one was able to inhibit PqsE enzyme activity intracellularly, as demonstrated by an *in vivo* enzyme assay. This finding encourages further development of the drug, Vorinostat, as it is already approved for use in humans and is able to bind intracellular PqsE in *P. aeruginosa*.

## Results and Discussion

### Competitive Fluorescence Polarization Screen

The initial stage of screening was performed using a fluorescence polarization (FP) assay testing 770 molecules of the SCREEN-WELL® FDA-approved drug library V2 from Enzo Life Science. This screen was designed to test whether the molecules could compete with the previously published BB562 fluorescent probe to bind in the PqsE active site ^21^. Each molecule in the library was tested at a single concentration of 250 µM with 2 µM purified PqsE, and a hit was determined to be any molecule that decreased fluorescence polarization by at least 25% when compared to the no-compound control (**Figure 1a**). Additional controls measuring fluorescence polarization of the probe alone, in the absence of purified PqsE were included on every plate. The initial single-dose screen yielded 28 molecules that were able to decrease fluorescence polarization by at least the target margin and moved into the next stage of the screen, a test for dose-dependent competitive binding (**Figure 1b**).

**Figure 1.**
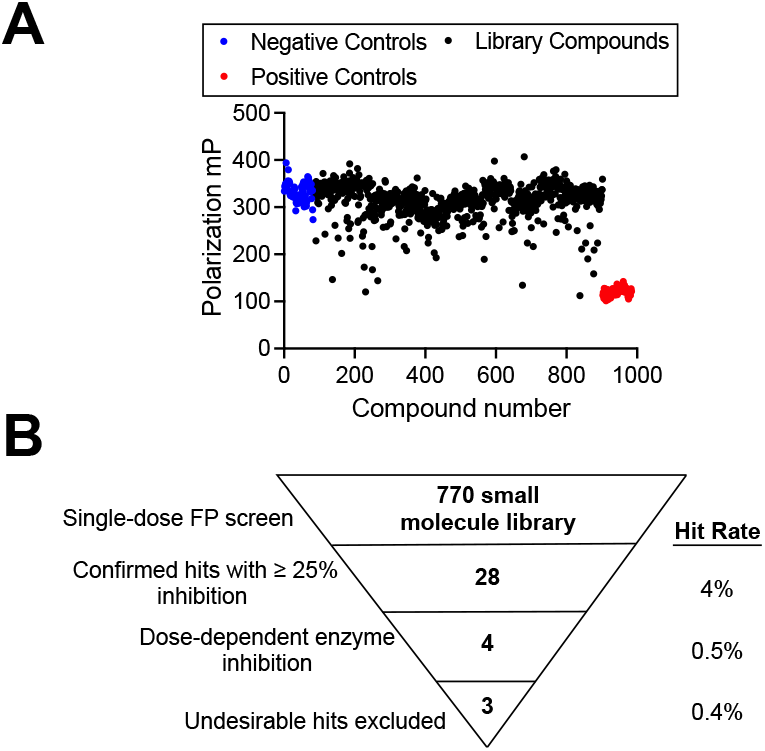
Competitive fluorescence polarization screen of FDA-approved small molecule library. A) Raw polarization values (in mP) obtained in initial signal-dose screen. Blue points represent the negative control with PqsE treated with DMSO, and red dots represent the positive control containing no protein. All black points represent library molecules and the fluorescence polarization measured for PqsE treated with that compound in the presence of the fluorescent probe, BB562. B) 770 molecules were screened at 250 µM and 28 molecules were identified to decrease fluorescence polarization by at least 25% compared to the DMSO-treated control wells (hit rate = 4%). Dose-response testing confirmed four molecules as hits, one of which was eliminated from further testing due to suspected off-target activity (hit rate = 0.4%).

The second stage of the screen took the 28 hit molecules from stage one and tested them in the same fluorescence polarization assay, but in a dilution series. The molecules were tested at a top concentration of 125 μM and in a two-fold dilution series to determine relative affinity of the hit molecules for PqsE. At this stage, the molecules were also tested for any potential spectral interference, as fluorescence polarization was measured for the dilution series of each test molecule plus the BB562 probe in the absence of PqsE. At this stage, a hit was determined to be any molecule that displayed a full inhibition curve (one exception being Trazodone, which did not reduce fluorescence polarization fully to baseline levels) (**Figure S1**). Notably, one molecule, (S)-Carbidopa, appeared to have significant dose-dependent inhibition, but upon repeated assays, was not a PqsE-binder (**Figure S1a**, P2-G7). Of the 28 molecules tested, four dose-dependent hits were identified and were purchased for further exploration.

The four dose-dependent hit molecules were then tested for their ability to inhibit an enzymatic activity of PqsE. PqsE is annotated as a thioesterase, but has broad hydrolase activity and is able to cleave a wide range of synthetic substrates *in vitro*. The hydrolysis of 4-methylumbelliferyl butyrate (MU-butyrate) has been used previously as a measure of PqsE enzyme activity ^21–23^. The four hit molecules were all able to dose-dependently inhibit PqsE enzyme activity with low to mid-micromolar potencies (IC_50_ values of 1.2 µM, 1.8 µM, 18 µM, and 180 µM for Vorinostat, Silver Sulfadiazine, Apomorphine, and Trazodone, respectively) (**Figure 2**). Upon further testing, Silver Sulfadiazine was eliminated from consideration due to its likely non-specific binding and destabilization of the protein. While exhibiting potent competition with the BB562 probe and inhibition of PqsE esterase activity, Silver Sulfadiazine was found to kill *P. aeruginosa* and thus, did not lend itself to further mechanistic testing *in vivo* ^24^.

**Figure 2.**
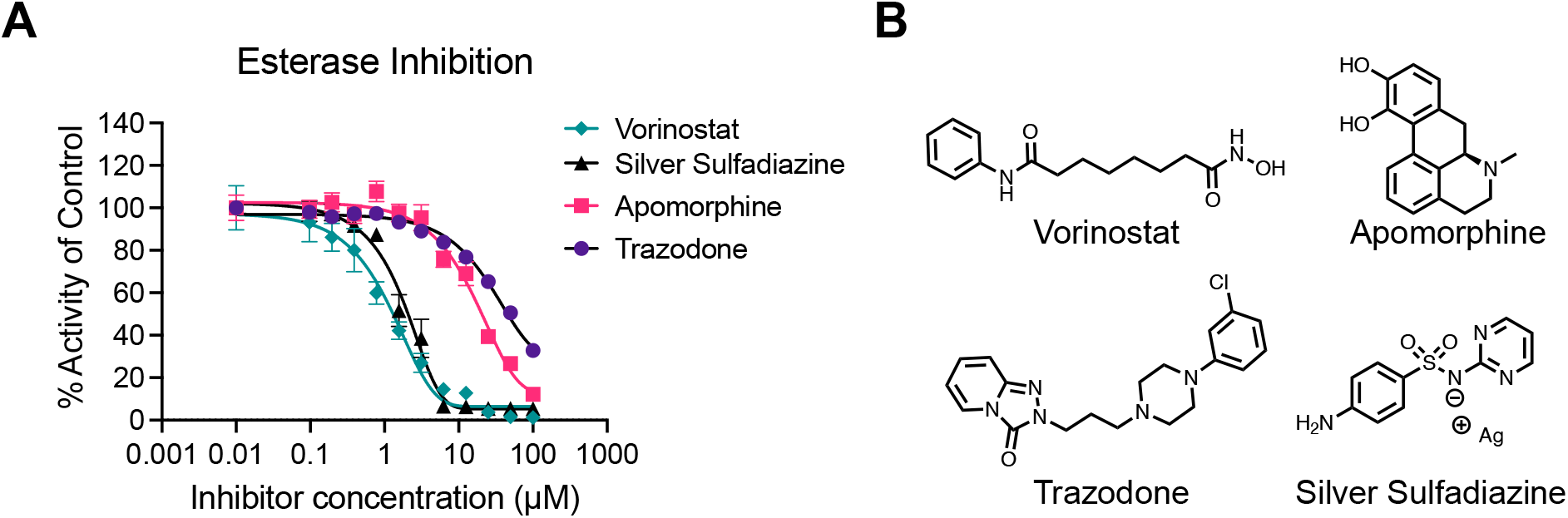
Inhibition of PqsE esterase activity A.) Dose-dependent inhibition of PqsE-catalyzed hydrolysis of MU-butyrate. % Activity is relative to the 0 µM, DMSO-treated protein and background fluorescence of the inhibitor dilution series in the absence of PqsE was subtracted from the data prior to normalization. Data points represent the average of technical triplicate values and error bars represent standard deviation. B.) Structures of the four confirmed dose-dependent screening hits.

### Inhibition of PqsE(WT) and variant with partially blocked active site

The hit molecules Vorinostat, Apomorphine, and Trazodone were analyzed further to understand the mechanism by which they decreased enzyme function. Silver Sulfadiazine was also tested in this manner (**Figure S2**), although the molecule was not subjected to further analysis. To this end, a previously established PqsE variant, PqsE(S285W) was used as a tool to determine whether the hit molecules were in fact directly binding in the active site of PqsE. The PqsE(S285W) variant has a partially blocked active site while retaining normal enzyme function compared to PqsE(WT) ^23^. Although the hydrolytic capacity of PqsE(S285W) is similar to that of the WT protein, this protein has a partially blocked active site, and exhibits approximately 10x lower affinity for the BB562 probe compared to PqsE(WT). If a hit molecule is binding in the active site, we would expect that molecule to have reduced potency against the PqsE(S285W) variant enzyme. Each of the hit molecules was tested in a multi-point dilution series against both PqsE(WT) and PqsE(S285W) in an esterase assay (**Figure 3**). Interestingly, only two molecules showed a decrease in potency against PqsE(S285W) (**Figure 3a** & **b**). The IC_50_ value of Vorinostat increased by 10-fold against PqsE(S285W), suggesting that this molecule binds in a way that is affected by the substitution of a serine (short sidechain, hydrogen bond donor/acceptor) with a tryptophan (bulky sidechain, no hydroxyl moiety for hydrogen bonding). Trazodone was also less potent against the PqsE(S285W) variant, however due to low potency and an incomplete inhibition curve, an IC_50_ value could not be determined. Apomorphine did not follow this trend (**Figure 3c**). Instead, the IC_50_ of Apomorphine was relatively unchanged for the PqsE(S285W) variant compared to PqsE(WT) (22 µM vs 26 µM, respectively). This suggests Apomorphine could be binding outside of the active site and allosterically affecting both binding of the BB562 probe and enzyme activity. This result is consistent with the structure of Apomorphine, which is likely too bulky to be able to bind in the active site of PqsE (**Figure 2b**).

**Figure 3.**
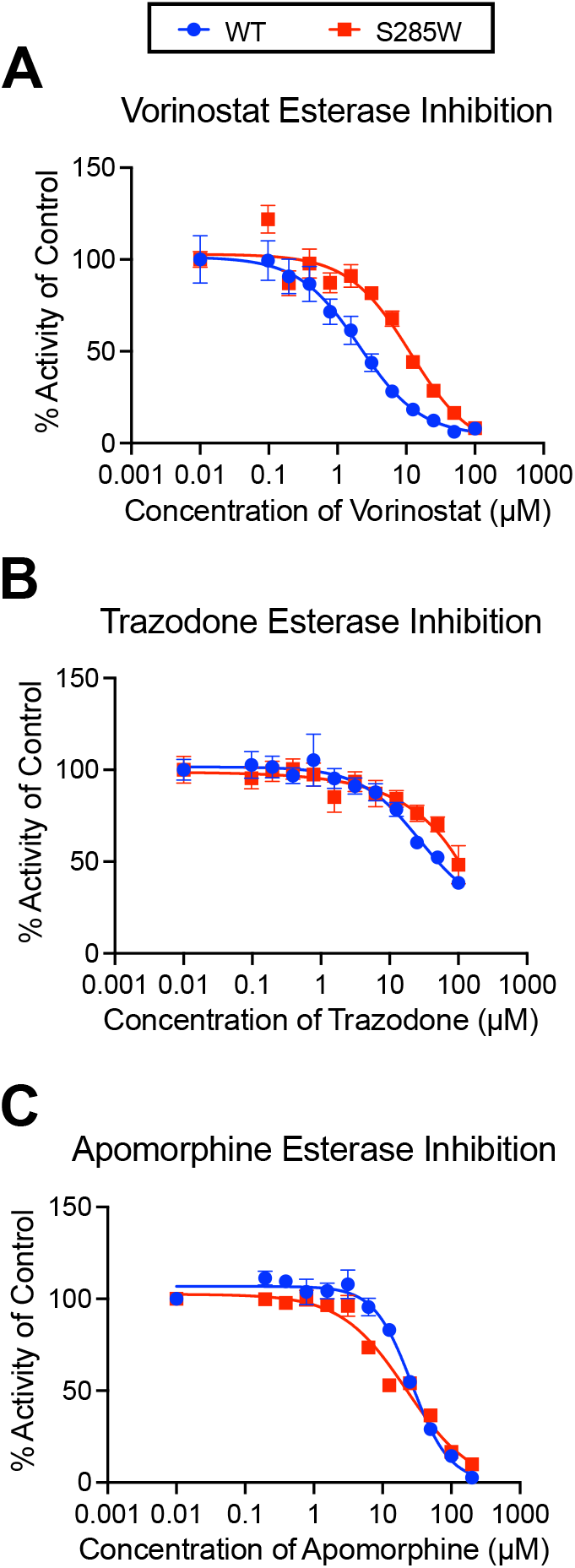
Inhibition of PqsE(WT) and PqsE(S285W) by screening hits. Esterase activity of PqsE(WT) (blue circles) and PqsE(S285W) (red squares) treated with varying concentrations of A) Vorinostat, B) Trazodone, and C) Apomorphine. % Activity is relative to the 0 µM, DMSO-treated protein and background fluorescence of the inhibitor dilution series in the absence of PqsE was subtracted from the data prior to normalization. Data points represent the average of technical triplicate values and error bars represent standard deviation.

### In vivo testing for PqsE-dependent inhibition of virulence phenotypes

The three hit molecules were also tested in *P. aeruginosa* cultures to determine their effects on QS-directed behaviors such as pyocyanin production. Pyocyanin production is an important virulence-related activity that is highly correlated to the ability of *P. aeruginosa* to infect a host ^25–27^. The ability to produce pyocyanin is dependent on the PqsE-RhlR protein-protein interaction. This protein-protein interaction, however, is completely independent of PqsE enzyme activity, as demonstrated by previously findings with the catalytically inactive PqsE(D73A) variant. The D73A mutation abolishes PqsE enzyme activity without affecting the ability of PqsE to form a complex with RhlR. It was found that *P. aeruginosa* strains expressing *pqsE(D73A)* produced the same levels of pyocyanin as strains expressing *pqsE(WT)*. Therefore, a molecule would only be able to inhibit pyocyanin production through a PqsE-dependent mechanism if that molecule is inhibiting the ability of PqsE to interact with RhlR. We constructed an assay to determine whether any inhibition of pyocyanin was specifically achieved via a PqsE-dependent mechanism. This was accomplished following the creation of a *P. aeruginosa* strain with an *azeB-lux* reporter ^28^. This reporter strain produces bioluminescence when the *azeB* gene is activated. In *P. aeruginosa*, the *azeB* gene is regulated by RhlR. However, unlike most RhlR-regulated genes, the interaction of PqsE with RhlR decreases expression of *azeB* ^28^. In this reporter strain, if the PqsE-RhlR complex is inhibited by a small molecule, we would expect to see a dose-dependent decrease in pyocyanin production as well as an increase in luminescence, meaning increased expression of *azeB*. This assay tests for specificity by confirming that the decrease in pyocyanin, a QS product that depends of the action of multiple biosynthetic enzymes, was accomplished through the direct inhibition of the PqsE-RhlR protein-protein interaction. If a molecule decreases pyocyanin production but does not increase luminescence, the pyocyanin decrease was not through PqsE and, therefore, is not on-target.

The three hit molecules, Vorinostat, Trazodone, and Apomorphine, were tested in this dual assay to look for on-target QS inhibition (**Figure 4**). Each molecule was tested at 50 µM and 100 µM in order to look for a dose-dependent trend. Each molecule was tested against two strains of *P. aeruginosa*: one that expressed *pqsE(WT)* and a Δ*pqsE* strain. This way, if any molecules had non-specific effects on luminescence (i.e. via inhibition of luciferase), this could be determined. All three molecules showed some slight effects on pyocyanin production, although none were statistically significant and mostly these effects were not dose-dependent. More importantly, the decreases in pyocyanin production were not accompanied by increases in *azeB* expression, suggesting that the mechanism by which pyocyanin production is being inhibited *in vivo* is not dependent on PqsE. Therefore, none of the hit molecules were shown to inhibit pyocyanin through inhibition of the PqsE-RhlR interaction in cell cultures. This finding was supported by an *in vitro* pull-down assay, in which all three compounds failed to inhibit complex formation between PqsE and RhlR (**Figure S3**). Interestingly, an increase in absorbance at 695 nm was observed in the Δ*pqsE* strain treated with Apomorphine (**Figure 4a**), but this was likely due to the slightly green color Apomorphine takes on upon oxidation ^29,30^.

**Figure 4.**
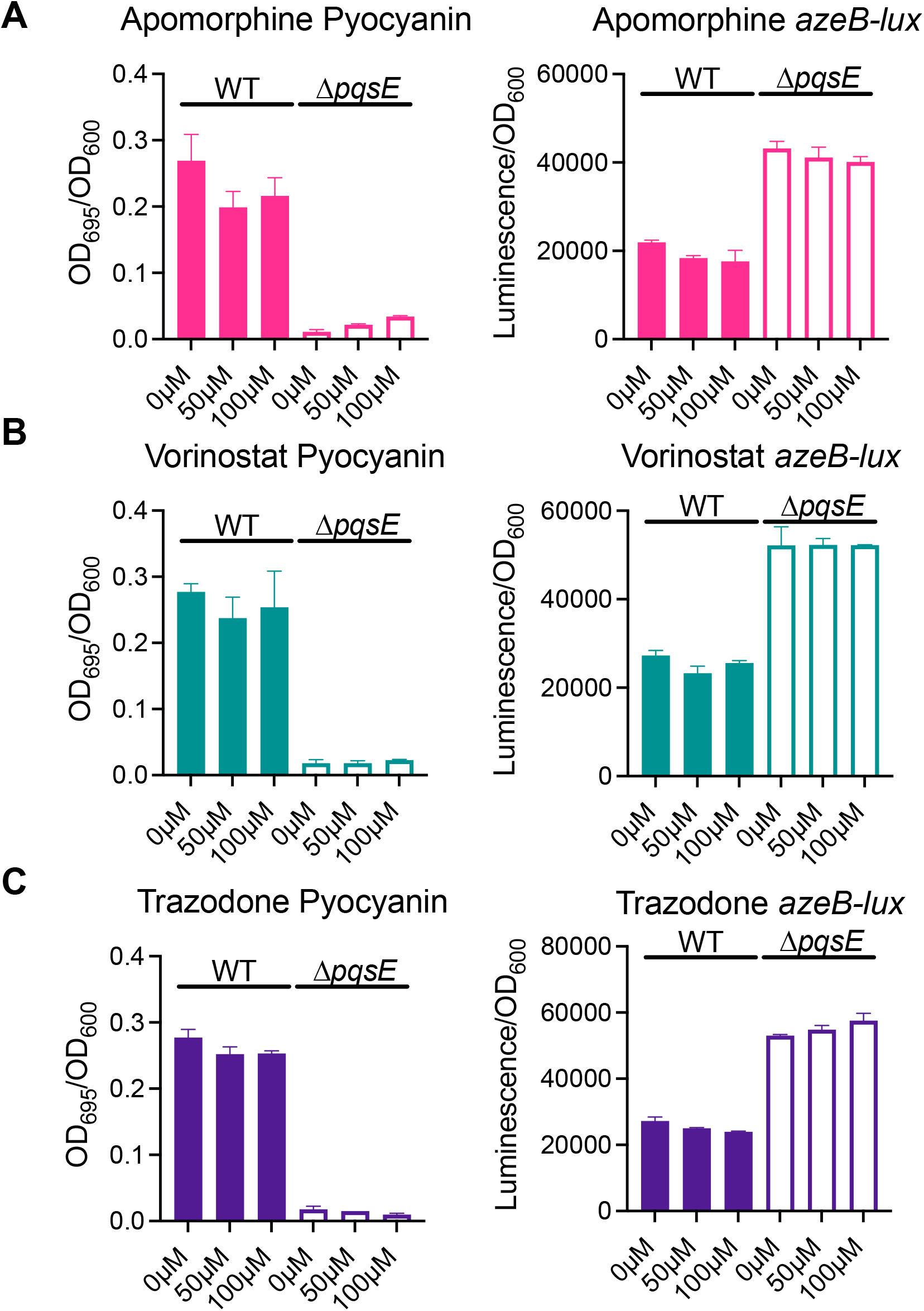
*In vivo* activity assays for pyocyanin production and *azeB* expression. Pyocyanin production is reported as OD_695_ of the culture supernatants divided by OD_600_ of the resuspended pellet. Activation of a transcriptional *azeB-luxCDABE* reporter was measured for the suspended pellet and reported as luminescence divided by OD_600_. The effect on both pyocyanin production and *azeB* expression was determined for A) Apomorphine, B) Vorinostat, and C) Trazodone in both a strain of *P. aeruginosa* expressing *pqsE(WT)* (filled bars) and a Δ*pqsE* strain (unfilled bars). Graphs show the average of two biological replicates. Error bars represent standard deviation.

### In vivo inhibition of PqsE catalytic function

Although the enzymatic function of PqsE is redundant to the biosynthetic pathway dedicated to producing the *Pseudomonas* Quinolone Signal (PQS), as demonstrated by the ability of Δ*pqsE P. aeruginosa* strains to produce PQS to wildtype levels^18,31^, this pathway can still be scrutinized to measure PqsE enzyme activity *in vivo*. PqsE performs a thioesterase function within the PQS synthetic pathway in order to convert 2-aminobenzoyl acetyl-CoA to 2-aminobenzoyl acetate (2-ABA) (**Figure 5a**). 2-ABA is inherently unstable, and to some extent, is decarboxylated to form a side-product, 2-aminoacetophenone (2-AA). 2-AA is a volatile molecule, and the characteristic fruity smell of *P. aeruginosa* cultures is attributed to its production ^32^. 2-AA has previously been used as a biomarker for the diagnosis of *P. aeruginosa* infections ^33^. We initially looked to 2-AA as a possible analyte for measuring enzymatic activity of PqsE in *P. aeruginosa*, however, when comparing 2-AA levels in the supernatants of a strain harboring catalytically inactive PqsE(D73A) compared to PqsE(WT), there was no significant difference in 2-AA levels as detected by comprehensive two-dimensional gas chromatography coupled with time-of-flight mass spectrometry (GC×GC-TOFMS). Further comparison of all analytes identified in the culture supernatants revealed that there was a statistically significant difference in concentrations of another molecule, acetophenone, between the WT strain and the *pqsE(D73A)* strain (**Figure 5b**). Acetophenone has also been explored as a biomarker for the diagnosis of *P. aeruginosa* infections, as it is produced to a significantly greater degree by *P. aeruginosa* compared to other species of bacteria ^34,35^. We reasoned that this could be a further breakdown product of 2-ABA, and therefore chose acetophenone as the analyte for measuring *in vivo* enzyme activity of PqsE.

**Figure 5.**
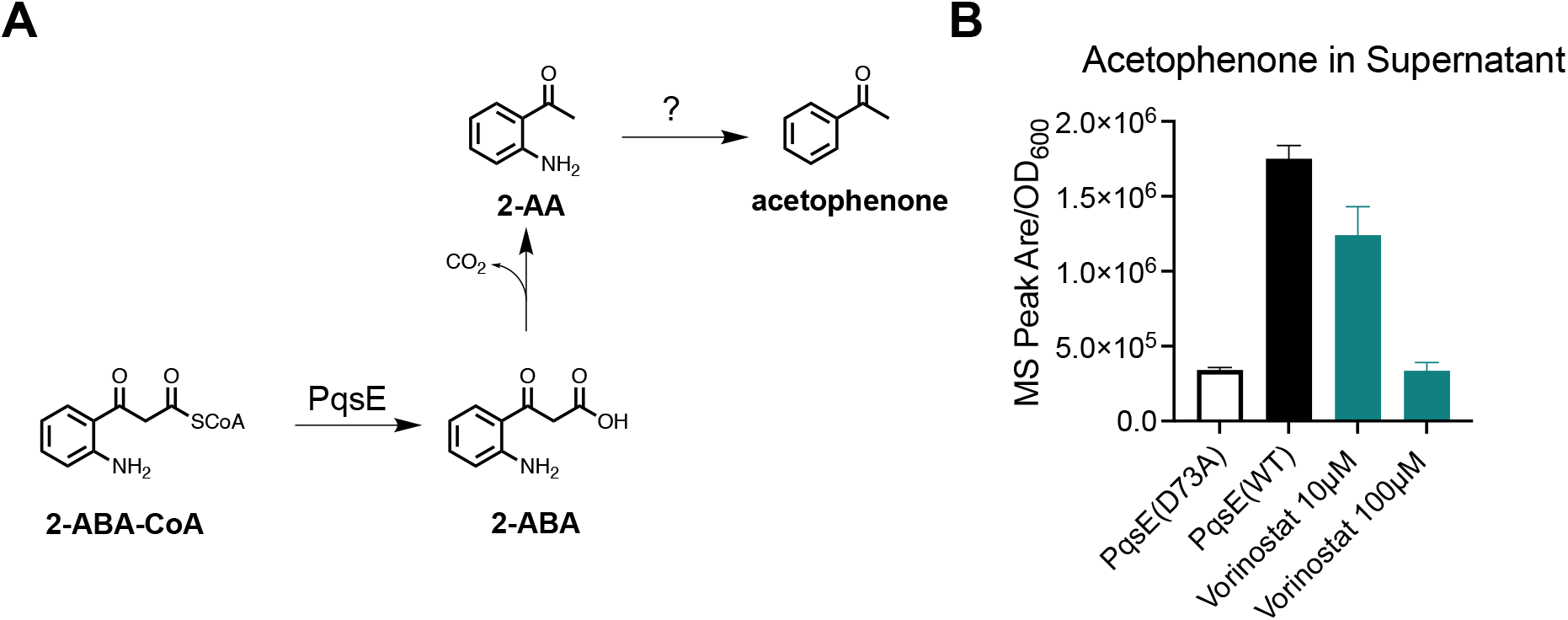
Assay for PqsE enzyme activity in *P. aeruginosa*. A) Reaction scheme for annotated PqsE catalytic function in PQS biosynthetic pathway and known by-products. The mechanism for PqsE-dependent production of acetophenone is uncertain, and is therefore annotated with “?”. B) Effect of Vorinostat on acetophenone production. Acetophenone levels detected in culture supernatants from *P. aeruginosa* strains grown for 8 hrs in the presence of specified concentrations of Vorinostat. All MS peak areas were first baseline-subtracted by the peak area detected in uninoculated LB and then normalized to OD_600_ of the suspended cultures to account for any differential growth rates. Data shown are the average of three biological replicates and error bars represent standard deviation.

To test whether the hit molecules, Vorinostat, Apomorphine, and Trazodone could inhibit PqsE enzyme activity in *P. aeruginosa*, cultures were grown in the presence of the inhibitors at 10 and 100 µM and acetophenone levels were measured in the culture supernatants after 8 hours of growth. A *P. aeruginosa* strain harboring *pqsE(D73A)* was grown as a control to determine the amount of acetophenone produced in the absence of PqsE catalytic activity. Compared to the *pqsE(D73A)* strain, the normalized peak area for acetophenone was approximately 5x greater for wildtype *P. aeruginosa*. When treated with 10 µM Vorinostat, there was a 30% decrease in acetophenone and when exposed to 100 µM Vorinostat, the amount of acetophenone detected dropped to the same level as detected in the *pqsE(D73A)* strain, suggesting full inhibition of PqsE catalytic activity (**Figure 5b**). Treating *P. aeruginosa* with either Apomorphine or Trazodone had no significant effect on acetophenone levels (**Figure S4**), suggesting that these molecules are either incapable of binding PqsE intracellularly, or they are not potent enough to have a measurable effect on PqsE enzyme activity *in vivo*. Either way, these molecules may require further synthetic optimization for use in *P. aeruginosa*.

## Experimental

### Strains, Media, and Chemicals

All experiments with *P. aeruginosa* used the UCBPP-PA14 strain as the parent strain (referred to as PA14). A list of all strains used in this study and their origin is included in the Supplementary Information (**Table S1**). Unless otherwise stated, all cultures were grown in Luria-Bertani (LB) broth (Difco). PqsE was recombinantly expressed and purified as described previously ^21,36^. The FDA-approved small molecule library was purchased from Enzo Life Sciences (BML-2843-0100, V.2.0) and screening hits were purchased individually from Enzo for follow-up testing.

### Fluorescence Polarization Screen

Methods were derived from Taylor *et. al* 2021^21^. Briefly, PqsE was diluted with assay buffer (50 mM Tricine, 0.01 % Triton X-100, pH 8.5) and added to the wells of an opaque 384-well plate at a final concentration of 2.0 μM. Each well had a final volume of 20 μL with the diluted protein accounting for 10 μL. Screen molecules diluted in DMSO were added to the wells at a final concentration of 250 μM and accounting for 0.5 μL of total well volume (2.5% DMSO final). The plate was incubated at room temperature for ∼10 minutes to allow protein-inhibitor complexes to form. The fluorescent probe, BB562, was diluted in assay buffer and added to the wells at a final concentration of 2.0 μM, accounting for 9.5 μL of the total well volume. The plate was then incubated at room temperature for 30 minutes before reading fluorescence polarization. Fluorescence polarization was measured in a Molecular Devices iD5 plate-reader with excitation and emission wavelengths of 485 and 530 nm, respectively. The Prism 9 software was used to generate all graphs and calculate EC_50_ values. In the dose-response FP assay, the inhibitor molecules underwent a two-fold dilution series before being added to the well plate. The top concentration in these assays was 125 μM.

### In vitro *Enzyme Assay*

To measure the enzyme activity of PqsE in the presence of an inhibitor, the general Esterase procedure was derived from Taylor et al. 2021^21^. Briefly, purified PqsE, diluted in assay buffer (50 mM Tricine, 0.01% Triton X-100, pH 8.5), was added to an opaque 384-well plate at a final concentration of 125 nM. The potential inhibitory molecules were diluted in DMSO and underwent a 2-fold dilution series to show the dose-dependent nature of their inhibition of enzyme function. Inhibitory molecules were added to the 384-well plate at a top concentration of 100 µM. The plate was incubated at RT for ∼5 minutes. MU-butyrate in assay buffer was added to the plate at a final concentration of 2 µM, with a final volume per well of 20 µL. The plate was incubated at RT for 20 min and then placed in a Molecular Devices iD5 plate-reader, where fluorescence of released 4-methylumbelliferone was measured (excitation: 360 nm, emission: 450 nm). To account for inhibitor fluorescence, control wells containing 2 µM MU-butyrate and the inhibitor dilution series were added to the wells in the absence of PqsE, still in a final volume of 20 µL per well. The Prism 9 software was used to generate inhibition curves and determine IC_50_ values. The same procedure was used for esterase assays involving PqsE(S285W).

### Dual Pyocyanin/azeB-lux Assay

The pyocyanin assay procedure was derived from Taylor et al. 2021^21^. Briefly, UCBPP PA14 strains of *P. aeruginosa* expressing *pqsE(WT)* or Δ*pqsE* with a chromosomally encoded *azeB-luxCDABE* fusion^28^ were grown overnight in LB media at 37 ° C with shaking. The next morning, the cultures were diluted 1,000x into 2 mL fresh LB, treated with various inhibitor concentrations (1% DMSO), and subsequently grown at 37 ° C with shaking. After 18 hours of growth, 1 mL of culture was pelleted by centrifugation at 14,000 rpm for 3 min. The supernatants were then collected, and the OD_695_ was measured in a UV/Vis spectrophotometer. The cell pellets were then resuspended in 1 mL PBS. The resuspended cells were loaded into the wells of a white clear-bottomed 96-well plate with 150 μL per well. The cell density (OD_600_) and luminescence were then measured in a Molecular Devices iD5 plate-reader. Pyocyanin production is reported as OD_695_ normalized to OD_600_ of the resuspended pellet and activation of *azeB-luxCDABE* is reported as luminescence/OD_600_.

### In vivo *Enzyme Assay*

*P. aeruginosa* PA14 strains expressing *pqsE(WT)* or *pqsE(D73A)* were grown overnight in LB at 37 ° C with shaking. The following morning, the cultures were diluted 100x into fresh LB treated with either DMSO or the test compounds in DMSO at the specified concentrations (1% DMSO final). The cultures were prepared in 6 mL total volume in 14 mL polypropylene round bottom tubes and grown at 37 °C with shaking at 200 rpm for 8 hr. Each condition was tested in triplicate, including three uninoculated LB blank samples with 1% DMSO. After 8 hr of growth in the presence of the test compounds, the cultures were removed from the incubator and 100 µL from each culture was transferred to a well of a clear bottom 96 well plate in order to measure cell density (OD_600_) in a Molecular Devices iD5 plate-reader. A low intensity orbital shake was completed prior to measurement to resuspend any settled cells. The remaining cultures were then pelleted by centrifugation at 4 ° C and 4000 rpm for 10 min. The supernatants were then filter sterilized through a 0.22 μm PES membrane into 20 mL headspace vials and then sealed with a PTFE/silicon cap. Immediately upon completing sample preparation, the headspace vials were transferred to a LECO Pegasus BTX 4D GC×GC-TOFMS instrument equipped with an LPAL autosampler for automated solid-phase microextraction (SPME) arrow headspace sampling and direct thermal desorption at the GC inlet. In the loading rack of the instrument, samples were at room temperature. Prior to injection, each headspace vial was agitated for 2 min at 35 ° C at 250 rpm. The full details of data collection and instrumental analysis are available in an extended methods section in the Supplementary Information.

## Conclusions

Due to its heightened antimicrobial resistance and a lack of viable treatment options, there is great interest in exploring new potential mechanisms for antibiotics to treat *P. aeruginosa* infections ^37^. Targeting virulence pathways in a way that would disarm but not kill bacteria is a strategy that is gaining traction in recent years, as the theory is, such an antibiotic would be less likely to spur the evolution of resistance mechanisms ^38,39^. Furthermore, there has been an increase in screening of FDA-approved small molecules, as this significantly expedites the drug discovery process ^40,41^. The Enzo FDA-approved molecule library ultimately yielded three molecules to explore in greater depth: Apomorphine, Vorinostat, and Trazodone. The hits were discovered using a multi-step screening process to identify molecules that competitively bind in the active site of PqsE. One surprising outcome of this screening strategy was that it not only identified competitive active site binders, but also identified one molecule, Apomorphine, that appears to bind outside of the active site and allosterically inhibit PqsE enzyme activity. These three hit molecules are all being explored further to determine their binding modes via structural techniques (primarily crystallography). We are particularly interested in solving a structure of Apomorphine bound to PqsE, as this would potentially be the first structure of a small molecule inhibitor bound to PqsE outside of the active site.

Upon *in vivo* testing, none of the three hit molecules could inhibit the PqsE-RhlR interaction specifically. Of the three molecules, there is compelling evidence that Vorinostat and Trazodone bind to the PqsE active site. Moving forward, co-crystal structures of these compounds bound to PqsE will allow the determination of points of interest for synthetic derivatization. The current most promising mechanism for PqsE-RhlR complex inhibition is for a small molecule to bind in the active site and induce PqsE to take on a configuration like that of the PqsE(E182W) variant. Since PqsE(E182W) is unable to bind to RhlR, inhibitor functional groups that reside near the E182 residue of PqsE will be flagged as the points of initial synthetic derivatization. Introducing strain near the E182 residue may induce the loop rearrangement that allosterically weakens the affinity of PqsE for RhlR.

Upon further *in vivo* analysis, only Vorinostat was found to inhibit PqsE enzyme activity in *P. aeruginosa*. This is particularly encouraging as previous screening efforts have produced molecules that potently inhibit PqsE enzyme activity *in vitro*, but were found to have no activity *in vivo* due to their probable membrane impermeability ^42^. This suggests that Vorinostat could serve as a starting point for further synthetic diversification, and would already possess the ability to cross the *P. aeruginosa* membrane and reach its target, PqsE, intracellularly. Although Apomorphine and Trazodone did not appear to inhibit enzyme activity of PqsE *in vivo*, these molecules were significantly less potent inhibitors *in vitro*, and their lack of intracellular activity could be due to their low potency. Apomorphine is still of great interest due to its structural diversity compared to previously identified PqsE inhibitors and potential new mode of action.

There is compelling evidence due to Apomorphine’s structure and the PqsE(S285W) esterase inhibition curve that supports the idea that the molecule is binding to an allosteric site on PqsE. Again, a co-crystal structure of Apomorphine bound to PqsE will reveal more about the mode of action of this inhibitor and whether it represents a viable pathway to explore for PqsE-RhlR complex inhibition. In addition to crystallography, we are exploring alternate ways to measure the ability of small molecules to trap PqsE in a configuration similar to that of PqsE(E182W). Some of these methods could be scaled to be able to carry out new high-throughput screens for such molecules.

## Author Contributions

H.A.J., M.J.B., N.Z., and A.L. conducted experiments and curated data. H.A.J., M.J.B., N.Z., A.L., K.A.P.U., and I.R.T. designed experiments and prepared the manuscript.

## Data Availability

The data supporting this article have been included as part of the ESI.

## Conflicts of Interest

There are no conflicts to declare.

## Acknowledgements

We thank members of both the Taylor and Perrault Uptmor groups for helpful advice and discussions. In particular, we thank Sarah Foster for assistance with developing the GC×GC-TOFMS methodology. We also thank Julie Valastyan and the Bassler Lab for lending the 2-aminoacetophenone standard that was used for GC-MS experiments. This work was supported by a Henry Dreyfus Teacher-Scholar Award for K.A.P.U.

## Notes

### Competing Interest Statement

The authors have declared no competing interest.

