## Supplemental Information for "Screening of FDA-approved small molecules to discover inhibitors of the *Pseudomonas aeruginosa* quorum-sensing enzyme, PqsE"

**This PDF file includes:**

Figure S1
Figure S2
Figure S3
Figure S4
Table S1
Extended methods

### Supplementary Figures

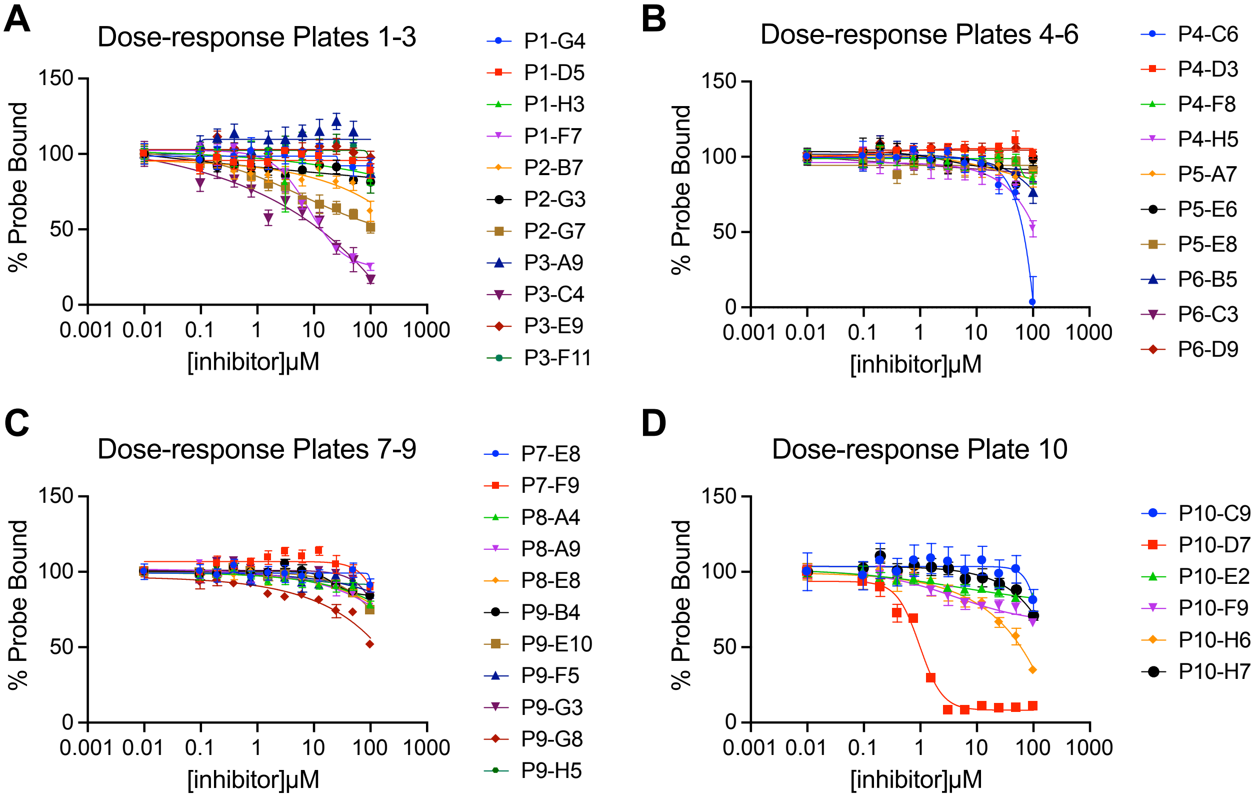


**Fig. S1.** Dose-response testing of hits from initial single-dose screen of FDA-approved molecules. Background fluorescence polarization (in the absence of PqsE) values were subtracted and measurements were normalized with polarization values at 0 μM competitor compound equal to 100% Probe Bound. Compounds that were re-tested in a dilution series were any initial hits that decreased fluorescence polarization by at least 25% compared to the DMSO control (28 hits) plus some select molecules that did not decrease fluorescence polarization. Values are plotted as the average of three technical replicates with error bars representing standard deviation. Inhibition curves were fit to the data in the Prism 9 software.

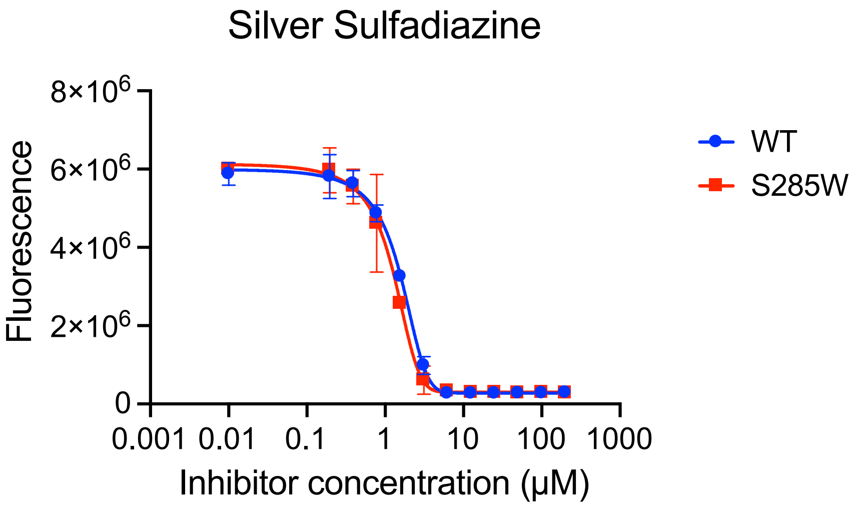


**Fig S2**: Assessment of Silver Sulfadiazine binding mode. Silver Sulfadiazine was tested for inhibition of the ability of both PqsE(WT) and PqsE(S285W) to hydrolyze the MU-butyrate ester substrate. $IC_{50}$ values against both the WT and variant protein with a partially blocked active site were nearly identical, suggesting that the binding mode of Silver Sulfadiazine does not rely on the active site serine. Values plotted are the average of technical triplicates with error bars representing standard deviation. The data shown are raw, unnormalized fluorescence values.

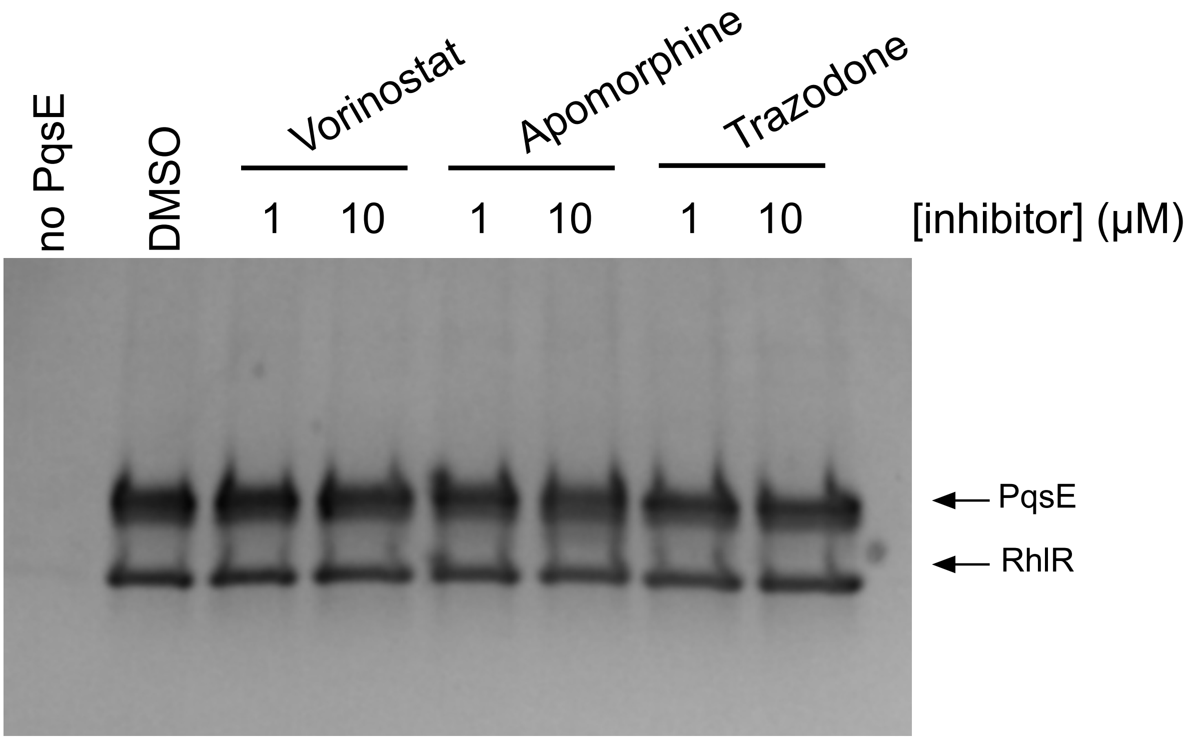


**Fig S3:** Effect of screening hits on in vitro PqsE-RhlR complex assembly. Purified PqsE with a 6xHis tag was incubated at 1 µM with the compounds Vorinostat, Apomorphine, and Trazodone at the specified concentrations. The PqsE:inhibitor complexes were subsequently incubated with lysate containing RhlR-mBTL and Nickel-coated resin. The resin was washed, proteins were eluted from the resin, and the eluates were subjected to SDS-PAGE. In the left-most lane, no PqsE was added and the RhlR-containing lysate was incubated with the Nickel resin to determine any non-specific binding. PqsE is ~34 kDa and RhlR is ~28 kDa, and the bands corresponding to each protein are labeled to the right of the gel image.

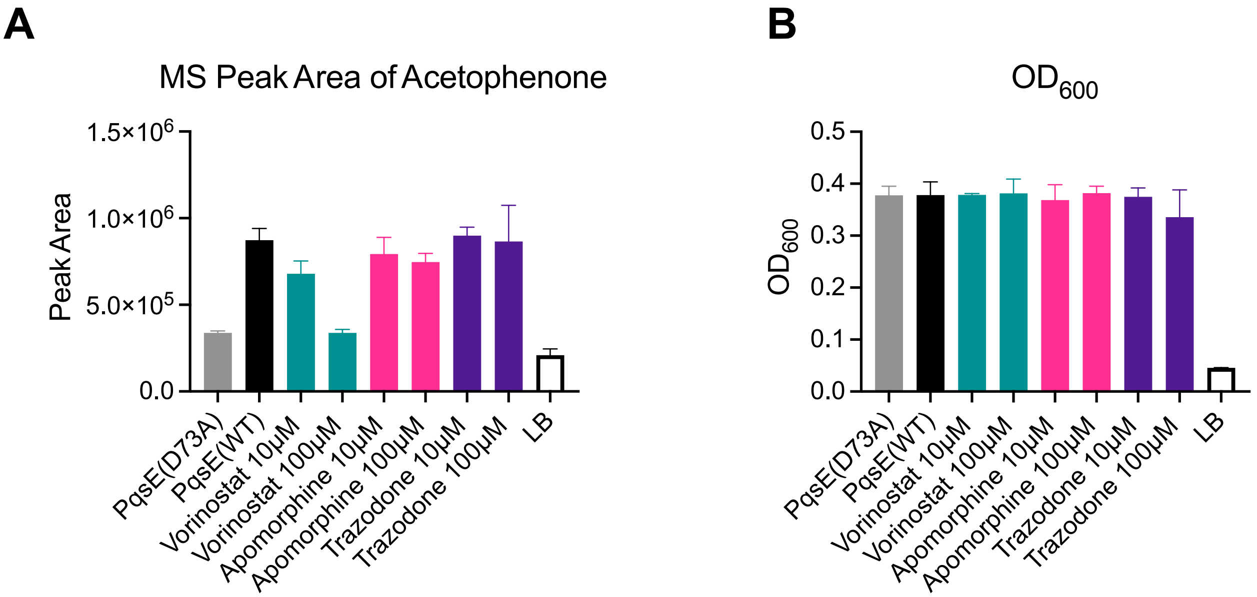


**Fig S4:** Cellular PqsE enzyme activity assay. A) MS peak area measured for acetophenone in culture supernatants. In all cases, including the PqsE(D73A) and PqsE(WT) controls, cultures were treated with 1% DMSO. Tubes of blank LB were incubated along with treated cultures to establish baseline measurements, which were subtracted from the data set for Vorinostat presented in Figure 5. B) The $OD_{600}$ was measured for each suspended culture at the end of the 8 hr growth period in the presence of the tested inhibitors. MS peak areas were normalized to $OD_{600}$ for the data presented in Figure 5. All data shown are the average of three biological replicates and error bars represent standard deviation.

**Supplementary Tables**

| **Strain** | **Description** | **Reference** |
|---|---|---|
| UCBPP-PA14 | PA14 *P. aeruginosa* Wildtype | Laboratory stock |
| SM776 | *E. coli* BL21 (DE3) pET28b-6xHis-pqsE(WT) | 36 |
| IT55 | *E. coli* BL21 (DE3) pET28b-6xHis-pqsE(S285W) | 21 |
| IT106 | PA14 *pqsE(D73A)* | 23 |
| IT245 | PA14 Δ*rhlI PazeB-luxCDABE* | This study |
| BT034 | PA14 Δ*rhlI*Δ*pqsE PazeB-luxCDABE* | This study |

**Table S1.** Strains used in this study.

### Extended Methods

*SPME Arrow Sampling*

Solid phase microextraction (SPME) arrow extraction was conducted with a 1.50 mm wide sleeve divinylbenzene/carbon wide range/polydimethylsiloxane (DVB/C-WR/PDMS) fiber (Restek Corporation, Bellefonte, PA, USA). Sampling was done on 20 mL headspace vials (Restek Corporation) containing 5 mL of supernatant. The fiber was chosen due to the range of analytes this sorbent can collect from headspace samples of a biological nature. Sample extraction and injection was performed using a LECO L-PAL3 Autosampler (LECO Corporation, St Joseph, MI, USA). After preparation, samples were transferred to the autosampler tray of the instrument. Sample incubation was performed for 2 min at 35 °C at 250 rpm. Sample agitation occurred at intervals of 5 s on followed by 2 s off. Sample extraction was performed for 5 min at 50 °C at 1000 rpm. The needle penetration depth was 40 mm into the sample vial and the penetration speed was 20 mm/s. Injection was performed to a depth of 40 mm at 10 mm/s with a desorb time of 2 min.

Prior to the sampling sequence, the SPME arrow fiber was conditioned at 270 °C for 40 min before the sequence and confirmed to be blank with a fiber blank injection. The SPME arrow fiber was reconditioned for 5 min prior to individual sample injection and for 2 min after injection at 270 °C. This reconditioning procedure was repeated between every sample, and an empty vial blank was included every nine samples.

*GC×GC-TOFMS Method*

The instrument used for analysis of *P. aeruginosa* supernatants was a Pegasus BTX GC×GC with a Paradigm Shift™ reverse fill-flush (RFF) flow modulator and dual channel detection using a flame ionization detector (FID) and a time-of-flight mass analyzer (LECO Corporation). The carrier gas was ultra-high purity helium (Airgas, Radnor, PA, USA) at a flow rate of 0.5 mL/min. The first-dimension column ($^1$D) was an Rxi-5MS column (20 m × 0.18 mm ID × 0.18 µm $d_f$, Restek Corporation). The second-dimension column ($^2$D) was an Rxi-17Sil MS (3.7 m × 0.25 mm ID × 0.25 µm $d_f$, Restek Corporation). The $^1$D flow rate was 0.5 mL/min and the $^2$D flow rate was 30 mL/min. The sample loop dimensions were 0.17 m × 0.53 mm ID resulting in a loop volume of 38 µL.

The modulation period was 4 s and the flush time was 158 ms throughout the duration of the run. This resulted in a flush factor of 1.73. The calculated flow to the TOFMS was 1.03 mL/min and the calculated flow to the FID was 37.18 mL/min. The inlet was operated in splitless mode for better detection of low-level analytes. The septum purge flow was 3 mL/min and the inlet purge time was 30 s with a purge flow of 20 mL/min and a total inlet flow of 50.5 mL/min.

The inlet temperature was 250 °C for the entire duration of the run. The initial temperature for the GC oven was 40 °C which was held for 2 min, then the oven was ramped at 5 °C/min until a target temperature of 230 °C was reached, with a final hold of 2 min, resulting in a runtime of 42 min. The transfer line was held at 345 °C and the ion source temperature was 300 °C. The TOFMS operated via electron impact (EI) ionization resulting in an acquisition rate of 100 scans/s and an extraction frequency of 30 kHz for the mass range of 30 – 550 *m/z*. An acquisition delay of 300 s was applied to mitigate solvent effects in the early part of the chromatographic run from saturating the MS signal.

The FID was set to 345 °C and operated at 100 Hz. The flow rate for hydrogen (ultra-high purity, Airgas) fuel was 45 mL/min. The flow rate for air (ultra zero purity, Airgas) was 450 mL/min. The flow rate for nitrogen (ultra-high purity, Airgas) makeup gas was 45 mL/min. The FID detector also had an acquisition delay of 300 Hz.

Data acquisition was performed for both detectors using LECO ChromaTOF software V5.59.02 with data processing V1.2.0.6 (LECO Corporation). GC×GC-TOFMS data was exported as .SMP files to a workstation computer with the same ChromaTOF processing software.

*Data Processing*

Samples were exported as .SMP files after the chromatographic run to a Network Attached Storage (NAS) system. Files were downloaded from the NAS to an offline data workstation and imported into a ChromaTOF database V5.59.02 with data processor V1.2.0.6 (LECO Corporation). This was done so that files could be then analyzed in ChromaTOF Tile® v1.3.50.0 (LECO Corporation). ChromaTOF Tile reviews raw GC×GC data to identify differences between samples or groups of samples based on class averages. A Fisher ratio test was performed on *pqsE(D73A)* and WT samples to identify molecules that showed significant differences in abundance between catalytically dead and active PqsE and thus could be effective in determining inhibitor effectiveness.

The parameters used for the statistical tests were a $^{1}$D tile size of 3 and a $^{2}$D tile size of 41 (auto-calculated in the software based on the height and width of a peak in ChromaTOF), $^{1}$D retention shift of 0 and a $^{2}$D retention shift of 0.01, a S/N threshold of 75, a total of 1 sample must exceed S/N threshold, 1 mass F-ratio to average, a minimum of 3 masses per tile, a minimum mass of 35 *m/z*, and a maximum mass of 550 *m/z*. A Fisher ratio threshold of 20 was applied.

After processing the data, a list of "hits" was generated which included the F-ratio value alongside the mean retention times in the first and second dimension, the quant mass, and a heatmap of relative amounts in each sample class average for each hit. The only "hit" reported was that of acetophenone, with a Fisher ration of 221.10. This hit was accepted based on comparison of its retention time and quant mass in ChromaTOF.

Following the discovery of a molecule effective for determining catalytic inhibition using the TOFMS signal, MS peaks for acetophenone were manually assigned in ChromaTOF to all samples to provide peak areas. MS peak areas for each sample were then transferred into Excel for preliminary analysis, normalization, and baseline subtraction. Lastly, Prism 9 software was used to generate all resulting graphs.
